# ROS signalling requires uninterrupted electron flow and is lost during ageing in flies

**DOI:** 10.1101/2021.08.18.456795

**Authors:** Charlotte Graham, Rhoda Stefanatos, Angeline E.H. Yek, Ruth V. Spriggs, Samantha H.Y. Loh, Alejandro Huerta Uribe, Tong Zhang, L. Miguel Martins, Oliver D.K. Maddocks, Filippo Scialo, Alberto Sanz

## Abstract

Mitochondrial Reactive Oxygen Species (mtROS) are cellular messengers essential for cellular homeostasis. In response to stress, reverse electron transport (RET) by respiratory complex I generates high levels of mtROS. Suppression of ROS produced via RET (ROS-RET) reduces survival under stress, while activation of ROS-RET extends lifespan in basal conditions. Here, we demonstrate that ROS-RET signalling requires increased electron entry and uninterrupted electron flow through the electron transport chain (ETC). We found that ROS-RET is abolished in old fruit flies where electron flux is reduced. Instead, mitochondria in aged flies produce consistently high levels of mtROS. Finally, we demonstrate that in young flies reduction of electron exit from the ETC, but not electron entry, phenocopies mtROS generation observed in old individuals. Our results define the mechanism by which ROS signalling is lost during ageing.

**Highlights:**

- ROS-RET signalling requires an uninterrupted flow of electrons through the ETC.
- ROS-RET signalling fails during ageing, with mitochondria producing persistently high levels of ROS.
- Interruption of ROS-RET signalling compromises stress adaptation in old flies.
- Reducing electron exit suppresses ROS-RET signalling and phenocopies ROS production observed in old mitochondria.

## 1. Introduction

A hallmark of ageing is the accumulation of damaged mitochondria that produce high levels of mitochondrial Reactive Oxygen Species (mtROS) [1]. The negative consequences of high ROS levels, i.e., loss of redox signalling and oxidative stress are well known [2], however, it is unclear how and why this occurs. In response to stress, *Drosophila* mitochondria produce ROS via Reverse Electron Transport (RET) (ROS-RET) [3]. ROS-RET occurs under conditions where both a highly reduced Coenzyme Q (CoQ) pool and elevated proton motive force (pmf) concur to allow RET from ubiquinol to CI [4]. Suppression of ROS-RET under stress prevents an adaptative transcriptional response and shortens survival in both fruit flies [3] and mice [5], while its stimulation in basal conditions extends lifespan [6]. This indicates the existence of a mitochondrial redox signalling pathway that regulates lifespan under both basal and stress conditions. This led us to hypothesize that ROS-RET signalling is affected by mitochondrial changes occurring during ageing which, in turn, limit stress adaptation in old individuals.

Here, we dissect the mechanism of ROS-RET signalling in detail. First, we combine high-resolution respirometry and metabolomic profiling to characterise how ROS-RET occurs *in vivo*. We show that ROS-RET is triggered by an increase in electron entry into the electron transport chain (ETC). Next, we demonstrate that ROS-RET signalling is lost during ageing and is replaced with sustained, high levels of ROS production. Finally, we dissect the alterations within the ETC that are responsible for the loss of ROS-RET signalling. We find that decreased electron exit suppresses ROS-RET signalling resulting in severe increases in ROS upstream of complex IV (CIV) which are unresponsive to stress.

## 2. Material and Methods

### 2.1 Fly husbandry

Wild type flies (white Dahomey, wDAH), RNA interference (RNAi) and GAL4 driver lines were maintained and collected as in [3]. Female flies 2-5 days old, unless otherwise stated, were used in all experiments. UAS-*levy*-RNAi (CG17280, 101523) and UAS-*ND*-75-RNAi (CG2286, 100733) were obtained from the Vienna Drosophila Resource Center (VDRC), while daughterless-GAL4 (daGAL4) was acquired from the Bloomington Drosophila Stock Centre. ETC inhibitors, rotenone and cyanide were added to the fly food at a final concentration of 600 µM and 18mM, respectively.

### 2.2 Thermal stress and lifespan studies

Thermal stress (TS) was induced by transferring flies from 25 to 32° for 3-4 hours. For all experiments except for lifespans, flies were used immediately after TS. For lifespan experiments, flies were exposed to TS three times per week for 4 hours in the presence or absence of rotenone and returned to 25°C (in food without rotenone) after TS.

### 2.3 Measurement of ROS in *Drosophila* brains

ROS were measured in individual fly brains as described in [3].

### 2.4 Measurement of mitochondrial oxygen consumption

Mitochondrial oxygen consumption was measured as described in [7].

### 2.5 Metabolomic Analysis by Liquid Chromatography-Mass Spectrometry (LC-MS)

Supernatants resulting from the homogenization of 20 fly heads were analysed by LC-MS as in [8]. LC-MS raw data were converted into mzML files using ProteoWizard. MZMine 2.10 was used for peak extraction and sample alignment and ANOVA was used to detect significantly altered metabolites (FDR < 0.05). These metabolites were analysed using Enrichment and Pathway Analysis with MetaboAnalyst [9].

### 2.6 Next-generation sequence data acquisition and analysis

RNA was extracted from 20 fly heads. Detailed experimental protocols and raw data are deposited in ArrayExpress under accession E-MTAB-7952. To select transcripts that were up-or down-regulated for subsequent analysis, we filtered transcripts whose Fold Change (FC) expression was above or below ±1.5, respectively, discarding those whose FDR was above 5%.

### 2.7 Statistical analysis

Data are shown as mean ± SEM and were analysed using GraphPad Prism 9 with either the unpaired Student’s t-test or One-way ANOVA with Dunnett’s post-test where appropriate. In addition, lifespan data were analysed using the Kaplan Meier Log-Rank Test. Cartoons and schematic diagrams were generated with the help of Biorender.

## 3. Results and Discussion

### 3.1 ROS-RET is activated by increasing electron flow into the ETC

We and others have shown that ROS-RET requires both an increase in the redox state of the CoQ pool and a sufficiently high pmf to occur [3, 4, 6]. Here, we dissected both the mechanism(s) that drive ROS-RET production and the downstream metabolic signals it triggers. The nature of the substrate oxidized by mitochondria determines where and how mtROS are generated [10, 11]. Accordingly, increased oxidation of substrates downstream of CI, such as succinate, fatty acids, or dimethylglycine, stimulates ROS-RET [10, 12, 13]. Furthermore, alterations in ROS levels reprogramme cellular metabolism in response to stress [14]. For example, a increase in cellular H_2_O_2_ activates the Pentose Phosphate Pathway (PPP) [15].

To understand how ROS-RET occurs, we exposed flies to TS and measured mitochondrial oxygen consumption. We found a significant increase in state 4 (phosphorylating) and state 3 (non-phosphorylating) respiration (Figure 1A), indicative of increased electron flow through the ETC. Accordingly, reducing electron flow by either depleting CI or complex II (CII) abolished ROS-RET [3]. Next, we performed unbiased metabolomics profiling on heads from flies exposed to TS (Figure S1A). To detect metabolic changes triggered by ROS-RET we fed flies with rotenone, which blocks RET from ubiquinol to CI [16]. Unsupervised Principal Component Analysis separated the three experimental groups (Figure 1B), suggesting that specific metabolic changes occured up-stream and down-stream of ROS-RET, with potential implications for signalling and stress adaptation. Pathway and Enrichment Analysis revealed alterations in amino acid levels associated with the initiation of ROS-RET (Supplementary Figure 1B-D). This is in line with previous observations describing that, under stress, glutamate or serine are rerouted to the ETC [17, 18]. Furthermore, manual inspection revealed changes in the levels of 11 metabolites with the potential to reduce ubiquinone (Figure 1C-D), supporting electron entry downstream of CI and CII as an important contributor to stimulation of ROS-RET.

**Figure 1.**
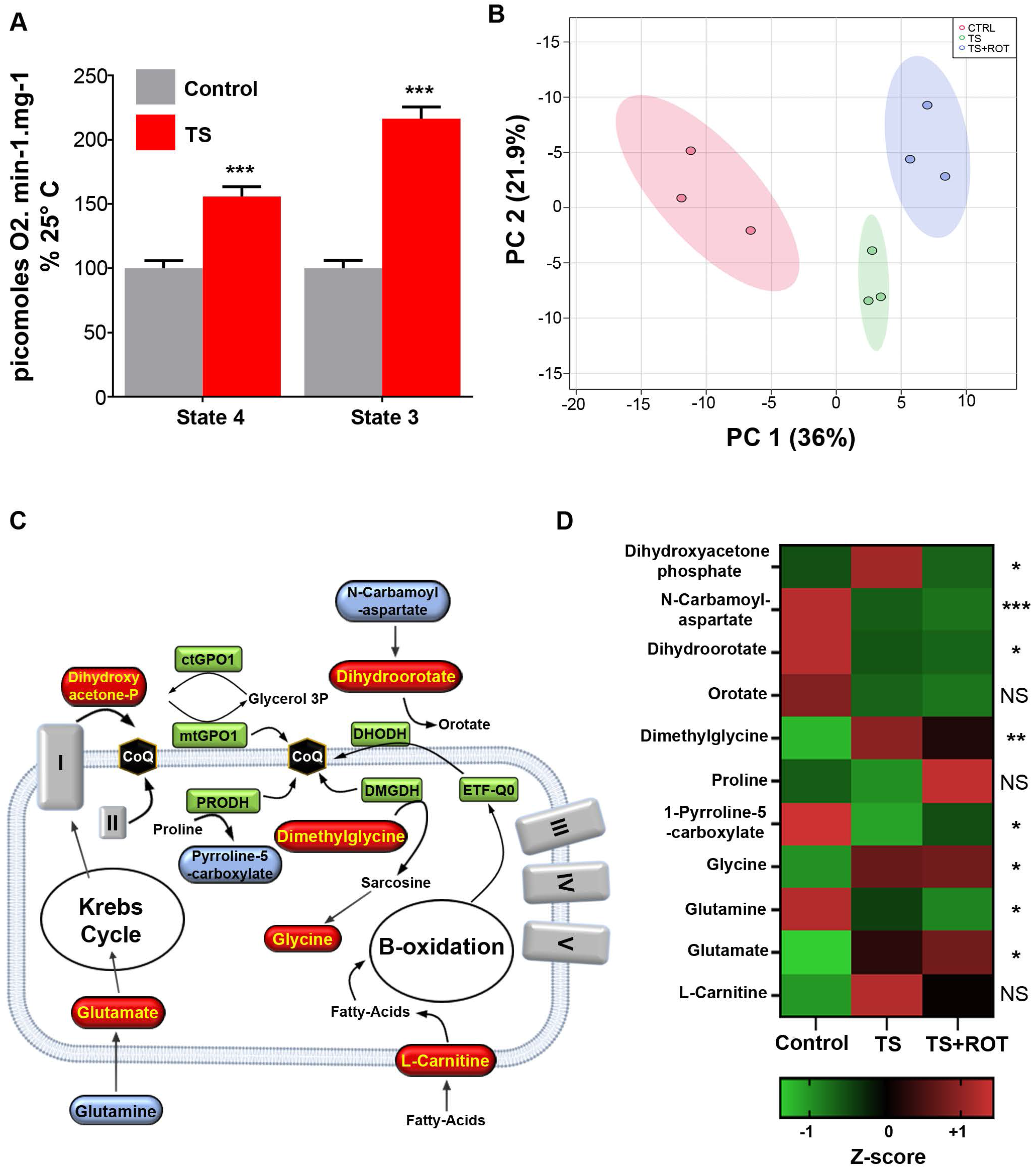
Thermal Stress (TS) increases electron flow through and downstream of CI. (A) Mitochondrial oxygen consumption measured in state 4 (without ADP) and state 3 (with ADP). Data are shown as mean ± SEM. (B) PCA analysis of the brain metabolome of Control flies and flies exposed to TS with (+ROT) and without rotenone. (C) Schematic representation highlighting key metabolites which show alterations in response to ROS-RET and their contribution to the reduction of ubiquinone to ubiquinol. Blue colouring indicates reduced levels while red indicates increased levels. cytoGPO1 = cytosolic Glycerol-3-phosphate dehydrogenase; mtGPO1 = mitochondrial Glycerol-3-phosphate dehydrogenase; PRODH = proline dehydrogenase; DHODH = Dihydroorotate dehydrogenase; DMGDH = Dimethylglycine dehydrogenase; ETF-QO = Electron-transferring-flavoprotein dehydrogenase. (D) Heat map of metabolites potentially contributing to the generation ROS-RET in response to TS. NS = not significant, *p<0.05, **p<0.01, ***p<0.001.

### 3.2 ROS-RET reroutes glycolytic intermediates to PPP but does not trigger a short-term transcriptomic response

We next investigated the metabolic changes initiated downstream of ROS-RET by selecting metabolites that were altered in response (up or down) to TS and were reverted by rotenone feeding (Figure S1A). Enrichment and Pathway Analysis showed PPP, glycolysis, purine, and glutathione metabolism as the main pathways modified by ROS-RET signalling (Supplementary Figure 2A-C). Our metabolomics analysis suggests that ROS-RET inhibits glycolysis, resulting in the redirection of glycolytic intermediates to PPP, allowing maintenance of NADPH levels (depleted during TS) and production of precursors for nucleotide biosynthesis (Figure 2B-D). Several observations support the notion that the ROS-RET activation of PPP is a general response to stress conserved across evolution. Firstly, NADPH was not detected in rotenone-fed flies (Supplementary Figure 2D), suggesting that ROS-RET is required for preventing depletion of NADPH via activation of PPP. NADPH depletion was not caused by increased oxidative stress since levels of GSH and GSSG were higher and lower, respectively, in the rotenone-fed group (Supplementary Figure 2D). Secondly, we found that upregulation of metabolites involved in the salvage purine biosynthesis pathway was suppressed when ROS-RET was inhibited with rotenone feeding (Figure 2D). This further supports the requirement of ROS-RET for a long-term transcriptomic anti-stress response. Accordingly, the expression of an alternative oxidase, which prevents ROS-RET, abolishes this response and shortens fly survival under TS [3] and mouse survival under mitochondrial stress [5]. Finally, a similar “glycolysis-to-PPP-rerouting” response has been reported in human keratinocytes and mouse macrophages exposed to increasing concentrations of H_2_O_2_ [15] or lipopolysaccharide (LPS) [19]. The latter is a particularly relevant example as increased ROS after LPS is a consequence of ROS-RET [20].

**Figure 2.**
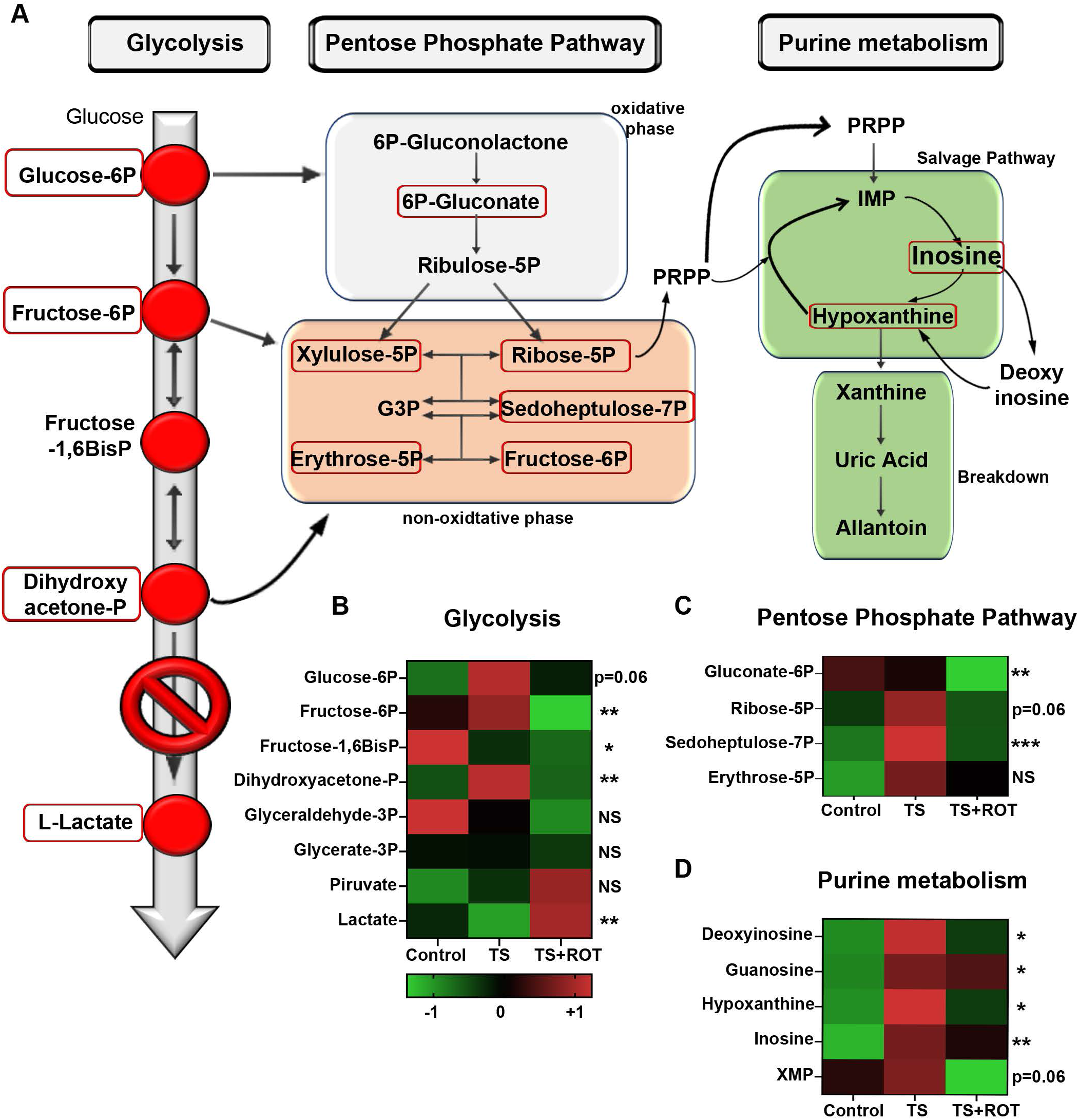
ROS-RET results in the redirection of glycolytic intermediates to the Pentose Phosphate Pathway (PPP) in order to produce NADPH and nucleotide precursors. (A) Schematic representation of the biological pathways affected by ROS-RET signalling. Metabolites significantly altered by ROS are indicated by red boxes. (C-D) Heat maps of the glycolytic, PPP and purine biosynthesis metabolites identified in the fly brain in control flies and flies exposed to TS with (TS+ROT) and without rotenone. NS = not significant, *p<0.05, **p<0.01, ***p<0.001.

We have shown that the long-term transcriptomic response to TS requires ROS-RET signalling [3]. Here, we analysed the brain transcriptome to investigate if manipulation of ROS-RET affects the short-term transcriptomic response to TS. As expected, we found upregulation of anti-stress pathways, such as heat-shock response (Sorensen, Nielsen et al. 2005). However, surprisingly, we also found that interruption of ROS-RET via rotenone did not cause any significant transcriptomic changes (Supplementary Figure 2E). These observations indicate that the ROS-RET is involved in the acute metabolic response to TS.

### 3.3 Ageing is characterized by consistently high levels of mtROS and the loss of ROS-RET signalling

High-resolution respirometry and metabolomic profiling show that ROS-RET relies on increased mitochondrial oxygen consumption. Accordingly, we and others have demonstrated that interrupting electron flow by depleting subunits of CI or CII prevents ROS-RET signalling [3, 6, 12, 21]. Since ageing is characterized by the reduction of mitochondrial respiration [6, 22], we anticipated that ROS-RET would be altered in old individuals. To test our hypothesis, we measured ROS levels in the brain of young (2-5 days), middle-aged (∼25 days), and old-aged flies (∼50 days). We found an accumulation of mtROS levels with age (Figure 3A). However, ROS-RET following TS was detected in middle-aged but not in old-aged flies (Figure 3B and 3C). Furthermore, rotenone treatment led to an increase in mitochondrial ROS levels in young but not old-aged flies (Supplementary Figure 3A,B). Since mitochondrial oxygen consumption is reduced in old-aged, but not middle-aged flies [6, 22], these findings support that the age-related reduction in mitochondrial function abolishes ROS-RET [6].

**Figure 3.**
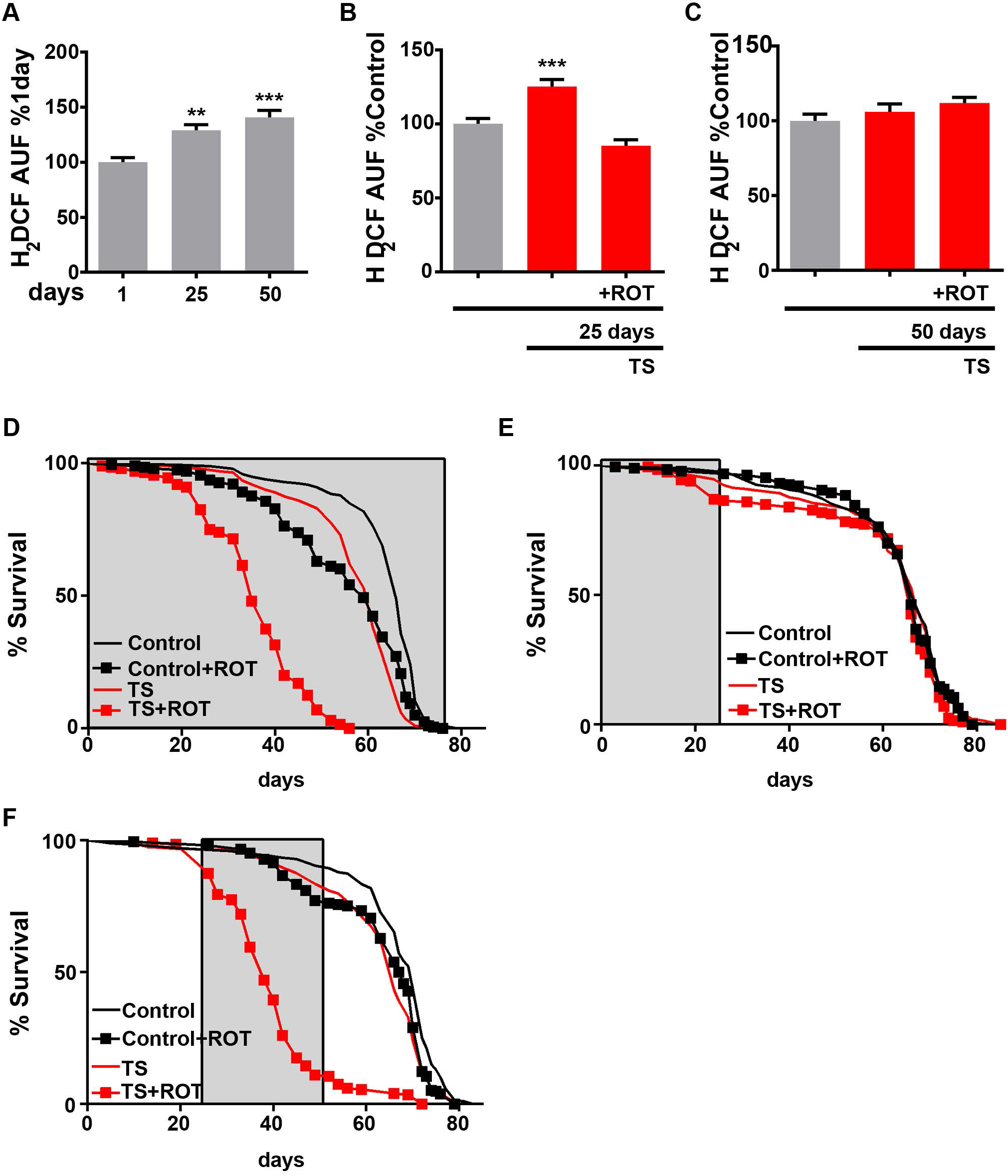
Old mitochondria produce continually high levels of ROS and are unable to activate ROS-RET signalling in response to stress. (A) ROS levels in brains of young (1 day), middle-aged (25 days) and old-aged flies (50 days). (B) ROS levels in brains of middle-aged flies exposed to TS in the presence (TS+ROT) or absence of rotenone. (C) ROS levels in brains of old aged flies exposed to TS with rotenone (TS+ROT) and without rotenone (TS). (D) Lifespan of control flies (black line), control flies fed with rotenone 3 times a week (black squares, Control+ROT), flies exposed to TS three times per week (red line, TS) and finally flies exposed to TS three times a week in the presence of rotenone (red squares, TS+ROT). Treatments were performed for the duration of fly lifespan. (E) As in D, but with treatments performed only between days 1 and 25 of fly lifespan. (F) Also as in D, but with treatments starting from day 25 to Day 50. (D-E) Grey shadow indicates duration of treatments. (A-C) Data are shown as mean ± SEM. ***p<0.001.

Next, we examined the physiological relevance of ROS-RET loss. For this, we exposed flies to TS for 4 hours, three times a week, in the absence or presence of rotenone in the food (Supplementary Figure 3C). Rotenone feeding severely shortened the lifespan of flies exposed to TS, whereas it minimally affected the lifespan of flies in basal conditions. However, this differential effect was only apparent when the flies were older than 20 days (Figure 3D). Accordingly, in an independent experiment where treatments (including rotenone) were stopped at day 25, no differences in lifespan were observed (Figure 3E). Conversely, rotenone treatment of flies between 25 and 50 days old dramatically shortened their survival (Figure 3F). These results indicate that the ability to produce ROS-RET progressively diminishes during ageing and that further suppression of ROS-RET in old individuals severely compromises survival.

### 3.4 Blocking mitochondrial electron exit mimics the mtROS profile observed in old individuals

Finally, we decided to study which alterations in the ETC were potentially responsible for the age-related increase in ROS levels observed in mitochondria from aged brains. First, we reduced electron entry by depleting CI subunit, *ND-75* (Supplementary Figure 4A). This caused a ∼50% decrease in mitochondrial respiration (Supplementary Figure 4B), analogous to the reduction observed in old flies [6, 22]. However, reduction of *ND-75* did not affect ROS levels in basal conditions but prevented ROS-RET under TS (Figure 4A-B). We also suppressed electron exit via depletion of CIV subunit, *levy* (Supplementary Figure 4C). This also decreased mitochondrial respiration by 50% (Supplementary Figure 4D). At steady state, *levy* knockdown increased ROS levels in the brain (Figure 4C). They remained high, with no observable increase caused by TS (Figure 4D-E). Finally, to test whether reducing CIV levels triggers ROS-RET, we fed flies with rotenone and measured ROS (Figure 4F). Since we did not observe a decrease, we could discard ROS-RET as the mechanism of ROS generation. To confirm the results of our genetic model, we used the specific CIV inhibitor, cyanide to acutely block CIV. Cyanide treatment inhibited respiration by over 50% (Supplementary Figure 4E) and increased ROS levels. As observed in the genetic model, rotenone treatment did not decrease ROS (Supplementary Figure 4F), discarding ROS-RET as the mechanism of ROS production.

**Figure 4.**
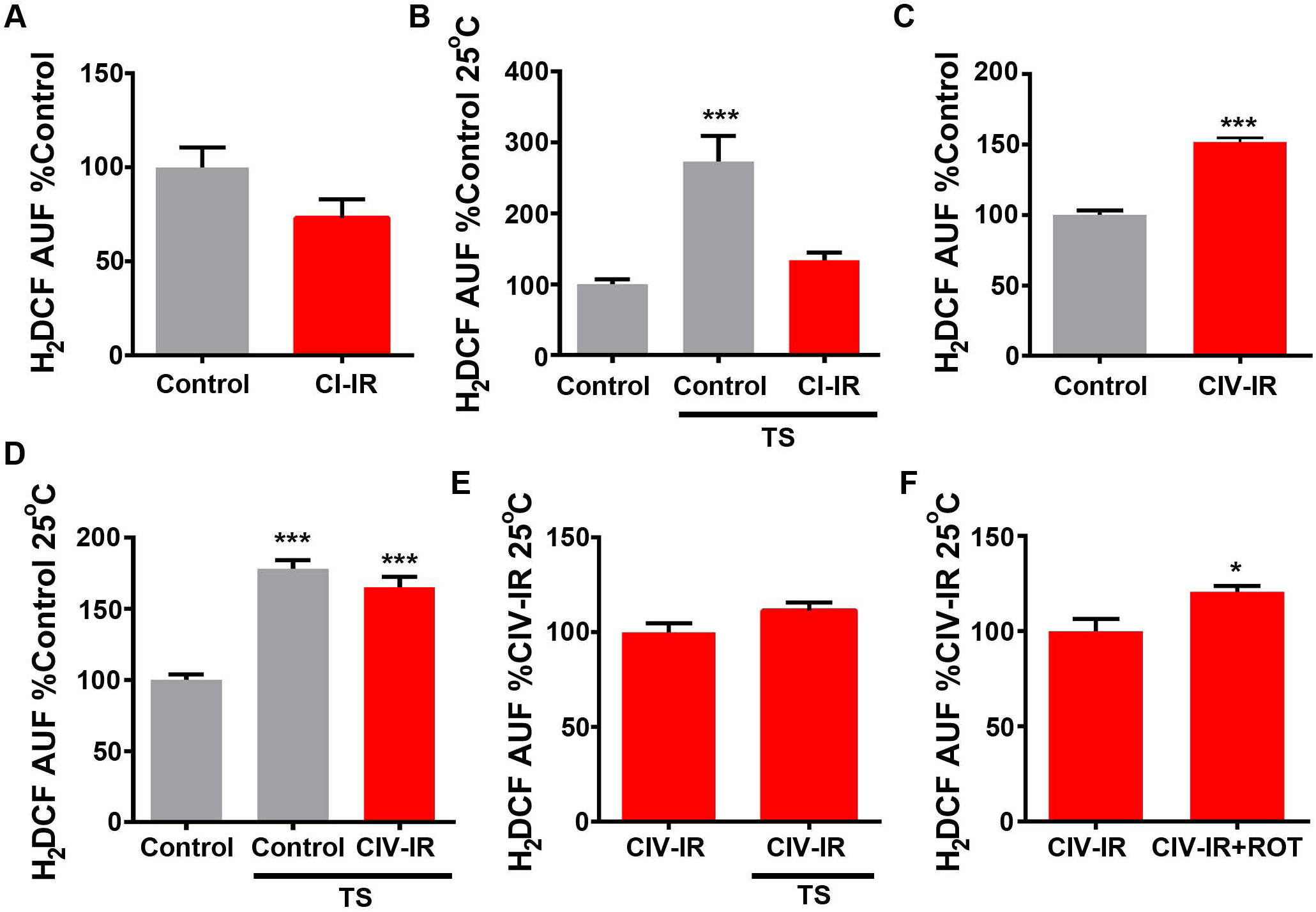
Depletion or inhibition of mitochondrial complex IV (CIV) mimics mitochondrial ROS production observed in old individuals. (A) ROS levels in brains of flies with reduced CI (CI-IR) and controls. (B) ROS levels in brains of flies with reduced CI (CI-IR) exposed to thermal stress and controls. (C) ROS levels in brains of flies with reduced CIV (CIV-IR) and controls. (B) ROS levels in brains of flies with reduced CIV (CIV-IR) exposed to thermal stress and controls. (E) Brain ROS levels in flies with depleted CIV levels (CIV-IR) at 25°C or under TS (32°C). (F) ROS levels in flies with reduced CIV (CIV-IR) in the presence or absence of rotenone (CIV+ROT). Data are shown as mean ± SEM. * P<0.05, ***p<0.001.

## 4. Conclusions

Our results indicate that ROS-RET occurs in young mitochondria when electron entry is increased through CI, CII and downstream mitochondrial dehydrogenases. Due to reductions in electron flow during ageing, the ETC starts to continually produce high levels of ROS but is unable to generate a ROS-RET signal in response to stress. Our results demonstrate that in young flies this is due to a reduction in electron exit. During ageing we propose that CI ceases to be the main ROS generator and is replaced by one or more generators inactive in young flies. The most likely candidate is CIII which has been shown to increase ROS when CIV is inhibited and has been described as the main generator of ROS in aged rat heart mitochondria [23-25]. Further research to confirm or discard this possibility is warranted.

## Abbreviations

H_2_O_2_: hydrogen peroxide
CI: complex I
CII: complex II
CIII: complex III
CIV: complex IV
CoQ: coenzyme-Q
CYA: Cyanide
ETC: electron transport chain
PPP: Pentose Phosphate Pathway
pmf: proton motive force
RET: reverse electron transport
mtROS: mitochondrial Reactive Oxygen Species
ROS-RET: ROS produced via Reverse Electron Transport
ROT: rotenone
TS: thermal stress.

## 5. Acknowledgements

This research was supported by a BBSRC grant (BB/R008167/2) and a Wellcome Senior Research Fellow (212241/A/18/Z) to AS, an MRC DTP studentship to CG, and a Henry Wellcome Postdoctoral Fellowship to RS (204715/Z/16/Z). RVS, SHYL and LMM are funded by the UK Medical Research Council, intramural project MC_UU_00025/3 (RG94521). AHU, ODKM and TZ were funded by a Cancer Research UK Career Development Fellowship (C53309/A19702).

## 6. Notes on authors contributions

CG performed ROS measurements, lifespan experiments and oxygraph analysis. RS assisted with lifespan and oxygraph experiments and edited the manuscript. AEHY performed oxygraphy analysis. RVS, SHYL and LMM generated the transcriptomics data. AHU, ODKM and TZ performed the metabolomic experiments. FS prepared samples for transcriptomics and metabolomics analyses. AS and FS designed and supervised the project, analysed data, assembled the figures and wrote the first version of the manuscript. All authors contribute to analyse and discuss the results and revised the manuscript.

## Figure Legends

**Supplementary Figure 1**. Linked to Figure 1. (A) Scheme depicts experimental design. We use three different groups: Control flies (that remained at 25°C), TS (flies exposed to 32°C for 4 hours) and TS+ROT (flies exposed to TS for 4 hours with rotenone in the food to prevent ROS-RET production). Metabolites triggering ROS-RET: Metabolites that were significantly changed in TS and TS+ROT and show the same trend in both groups. Metabolites regulated by ROS-RET: metabolites that were significantly changed in TS and TS+ROT but trends were opposite. (B) Enrichment analysis of metabolites stimulating ROS-RET. (C-D) Pathway analysis of metabolites triggering ROS-RET.

**Supplementary Figure 2**. Linked to Figure 2. (A) Enrichment analysis of metabolites altered by ROS-RET signalling. (B-C) Pathway analysis of metabolites altered by ROS-RET signalling. (D) Heat maps showing metabolites involved in glutathione metabolism identified in the fly brain of controls and groups exposed to Thermal Stress (TS) with (TS+ROT) or without rotenone (TS). NS = not significant, *p<0.05, **p<0.01. (E) Volcano plot showing differences in gene expression between flies exposed to TS in the absence or presence of rotenone (TS+ROT). Only one transcript was significantly upregulated in response to ROS-RET (indicated in red).

**Supplementary Figure 3**. Linked to Figure 3. (A) ROS levels in brains of young flies fed with rotenone (ROT). (B) ROS levels in brains of middle-aged flies fed with rotenone (ROT). (C) Schematic representation of the experimental design showing experimental conditions. 4 groups of flies were used, two groups were exposed to TS for 4 hours three times per week in either the absence or presence of rotenone. An additional control group at 25°C was fed with rotenone 4 hours three times per week. Treatments were implemented for either the duration of lifespan (see Panel 3D), from day 1 to day 25 (see Panel 3E), or from day 25 to day 50 (see Panel F). (A-B) Data are shown as mean ± SEM. ***p<0.001.

**Supplementary Figure 4**. Linked to Figure 4. (A) Schematic illustrating the strategies used to reduce electron entry into the ETC. (B) Mitochondrial oxygen consumption in homogenates from CI-depleted flies and controls. (C) Schematic diagram with the strategies used to reduce electron exit from the ETC. (D) Mitochondrial oxygen consumption in homogenates from CIV-depleted flies and controls. (E) Mitochondrial oxygen consumption in head homogenates from flies fed with cyanide (CYA) and controls. (F) ROS levels in brains from control flies, flies fed with cyanide (CYA) and flies fed with cyanide and rotenone (CYA+ROT). Data are shown as mean ± SEM. **p<0.01, ***p<0.001.

## Notes

### Competing Interest Statement

The authors have declared no competing interest.

