## Supplementary material for "ROS signalling requires uninterrupted electron flow and is lost during ageing in flies": Figure S1

A

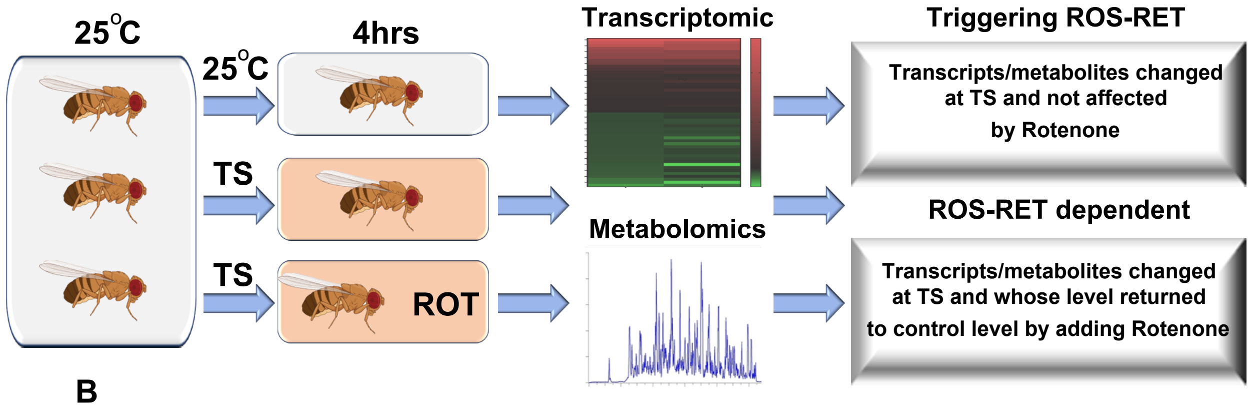


B

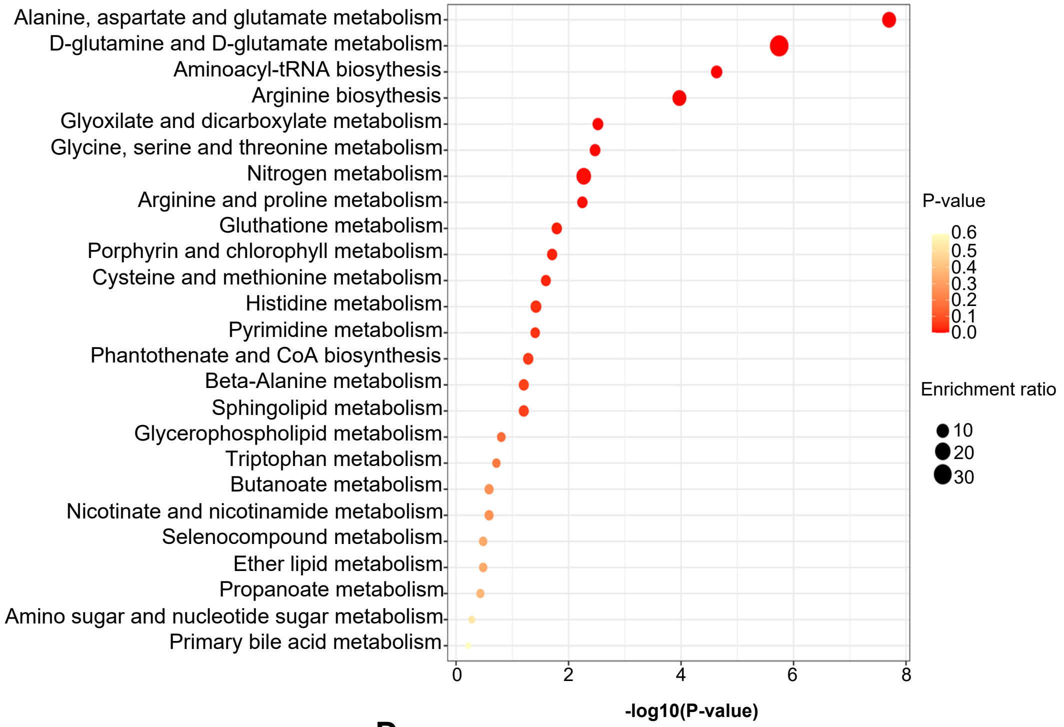


C

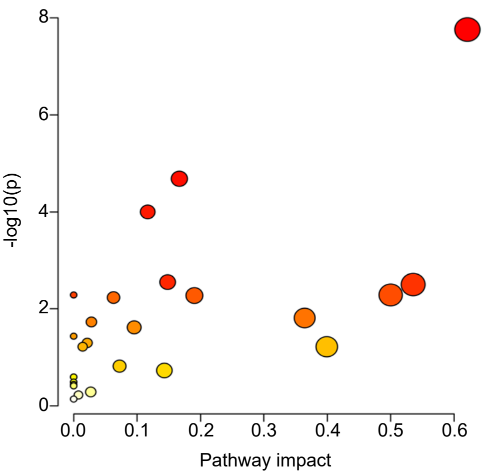


D

| | | Total | Expected | Hits | p-value |
|---|---|---|---|---|---|
| 1 | Aminoacyl-tRNA biosynthesis | 48 | 0.92903 | 7 | 2.07E-05 |
| 2 | Arginine biosynthesis | 14 | 0.27097 | 4 | 1.00E-04 |
| 3 | Glyoxylate and dicarboxylate metabolism | 32 | 0.61935 | 4 | 0.002816 |
| 4 | Glycine, serine and threonine metabolism | 33 | 0.63871 | 4 | 0.003161 |
| 5 | Nitrogen metabolism | 6 | 0.11613 | 2 | 0.005178 |
| 6 | D-Glutamine and D-Glutamate metabolism | 6 | 0.11613 | 2 | 0.005178 |
| 7 | Arginine and proline metabolism | 38 | 0.73548 | 4 | 0.00533 |
| 8 | Pyrimidine metabolism | 39 | 0.75484 | 4 | 0.00586 |
| 9 | Gluthatione metabolism | 28 | 0.54194 | 3 | 0.01547 |
| 10 | Porphyrin and chlorophyll metabolism | 30 | 0.58065 | 3 | 0.01868 |
| 11 | Cysteine and methionine metabolism | 33 | 0.63871 | 3 | 0.024141 |
| 12 | Histidine metabolism | 16 | 0.30968 | 2 | 0.036734 |

**Supplementary Figure 1**
