## Supplementary material for "ROS signalling requires uninterrupted electron flow and is lost during ageing in flies": Figure S2

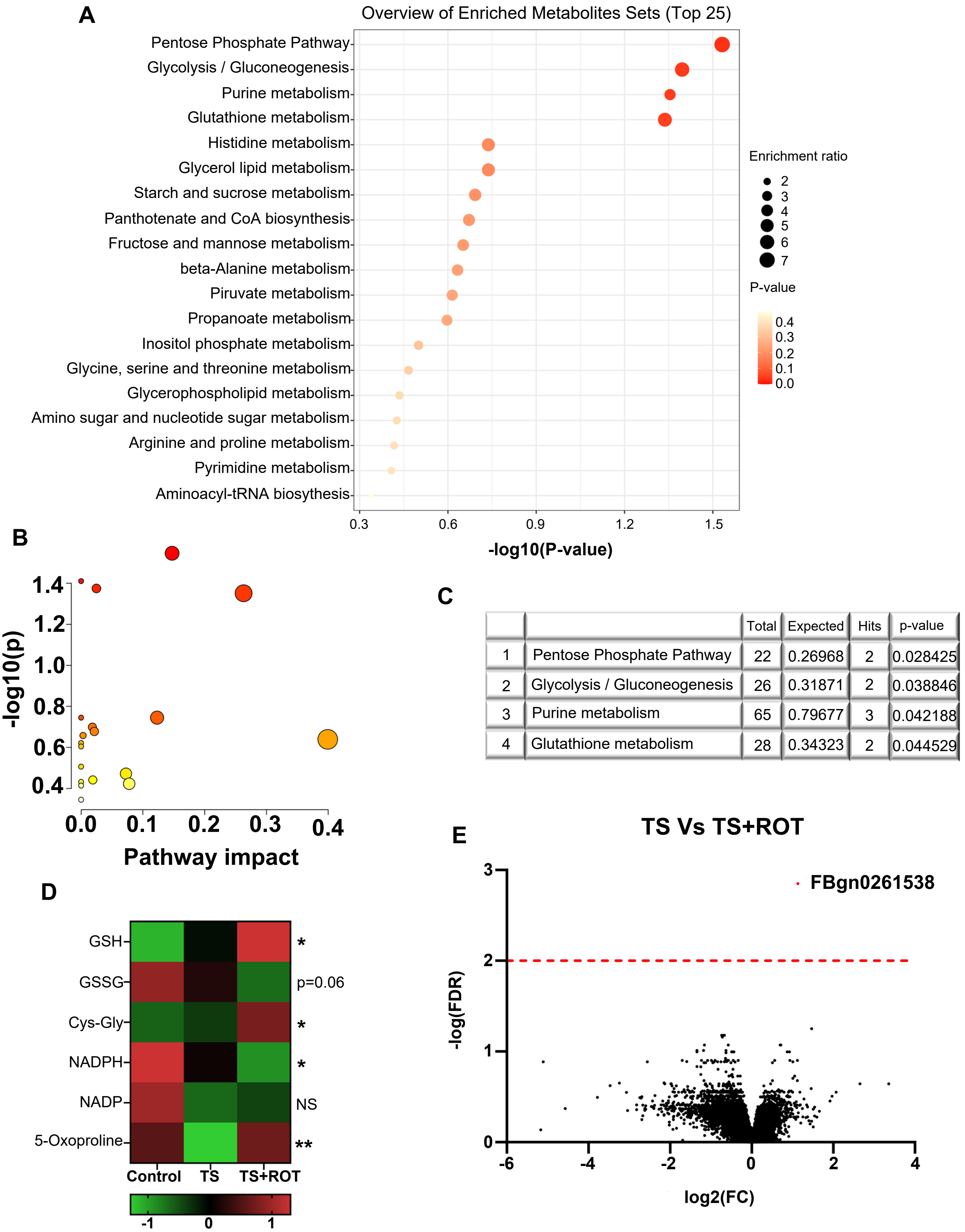


| | | Total | Expected | Hits | p-value |
|---|---|---|---|---|---|
| 1 | Pentose Phosphate Pathway | 22 | 0.26968 | 2 | 0.028425 |
| 2 | Glycolysis / Gluconeogenesis | 26 | 0.31871 | 2 | 0.038846 |
| 3 | Purine metabolism | 65 | 0.79677 | 3 | 0.042188 |
| 4 | Glutathione metabolism | 28 | 0.34323 | 2 | 0.044529 |

**Supplementary Figure 2**
