## Supplementary figures and images for "ROS signalling requires uninterrupted electron flow and is lost during ageing in flies"

### Figure S3

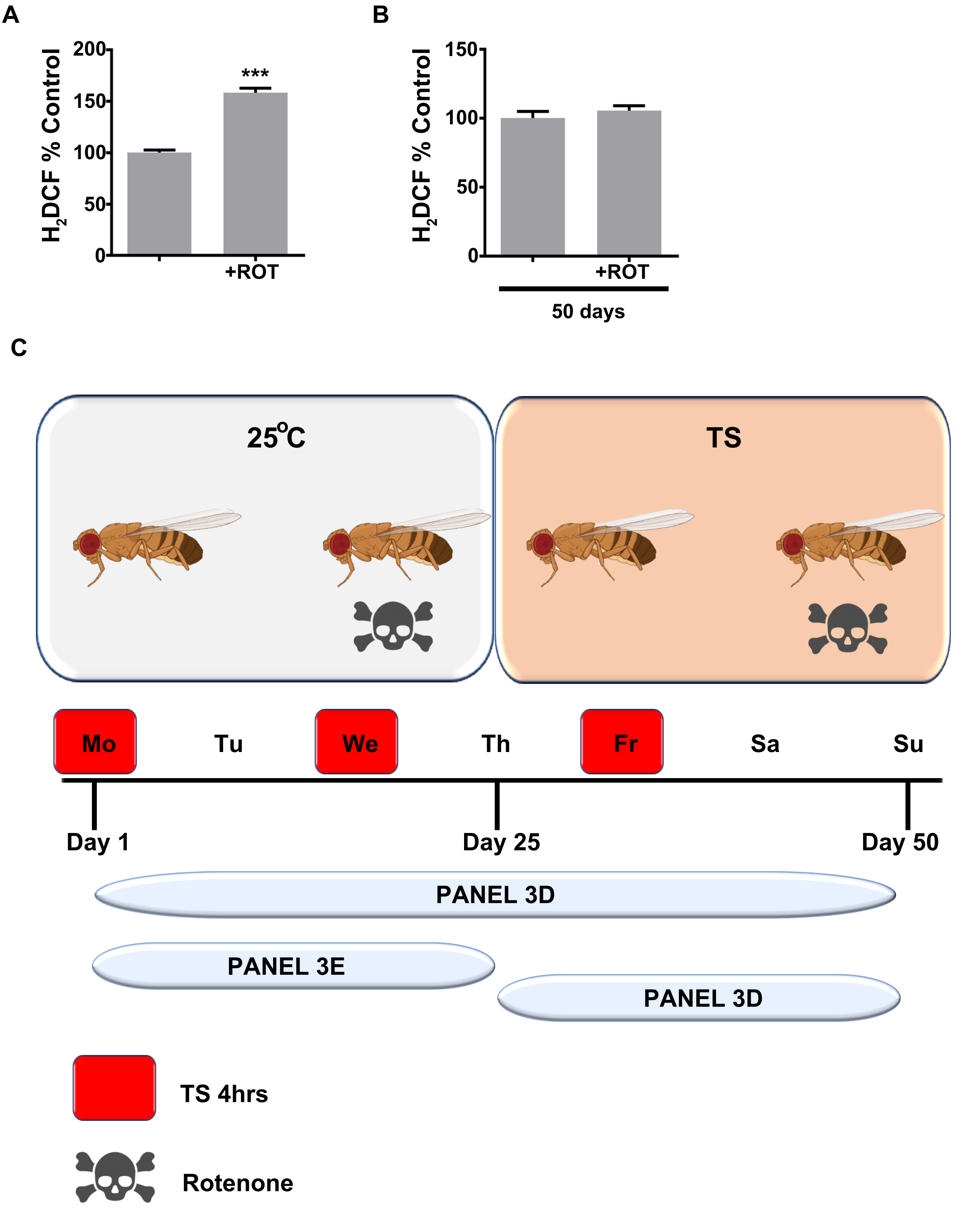


Supplementary Figure 3

### Figure S4

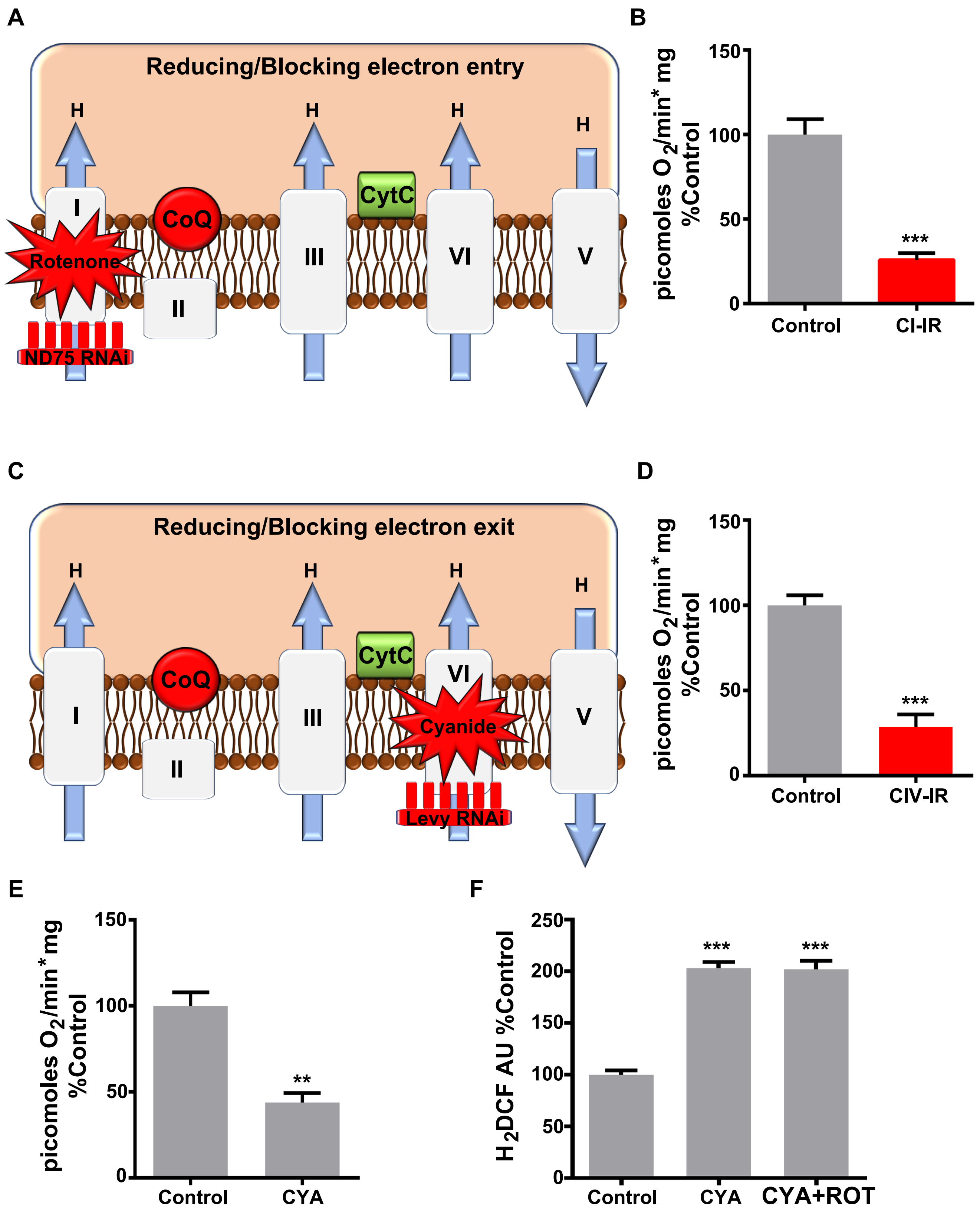


Supplementary Figure 4
